# Automated identification of ataxia and convulsions in hyperexcitable *Shudderer* mutants of the *Drosophila* sodium channel gene *paralytic*

**DOI:** 10.1101/2025.10.22.683992

**Authors:** Victoria Hand, Atulya Iyengar

## Abstract

*Drosophila* carrying the *Shudderer* (*Shu*) allele of the voltage-gated Na^+^ gene *paralytic* display spontaneous convulsions and immobilization phenotypes that are exacerbated by high temperature. To automate identification of these aberrant behaviors in *Shu* mutants, we trained a machine-learning classifier on a manually annotated dataset. The system reliably classified walking activity and immobilization periods, while uncoordinated movement events were detected with moderate sensitivity and high specificity. We then characterized the behavioral repertoire of *Shu* mutants and wild-type flies over a temperature-ramp protocol (20-40 °C) using the classifier. Our developments facilitate quantitative studies of environmental or genetic factors that alter behaviors characteristic of *Drosophila* models of neurological disease.

## Description

In *Drosophila*, disruptions in ion channel genes that tune membrane excitability or in genes encoding synaptic transmission machinery often lead to characteristic motor phenotypes such as paralysis, tremors, spasms or convulsions (Ganetzky & Wu, 1982, 1985; Siddiqi & Benzer, 1976; Suzuki et al., 1971). Neurogenetic analysis of flies harboring mutations in these genes have facilitated identification of key factors (genetic or environmental) contributing to the emergence of neurological phenotypes in humans carrying mutations in orthologous genes.

Automated video tracking of fly behavior in open arenas represents a convenient and versatile approach to monitor *Drosophila* motor phenotypes in a high-throughput fashion with minimal variability due to observer differences. Compared to manual observation approaches such as climbing assays (Benzer, 1967; Gargano et al., 2005), or water bath-based high-temperature sensitivity assays (Ganetzky & Wu, 1982), video tracking can quantitatively describe a larger set of locomotion parameters from a single observation (see Wang et al., 1997, and Martin, 2004 for early examples). More sophisticated machine learning approaches applied to video tracking data facilitate identification of complex behavioral sequences such as courtship-related song production, aggression-related lunges, sleep-related posture changes and grooming sequences (Berman et al., 2016; McKenzie-Smith et al., 2025; Ning et al., 2022; Qiao et al., 2018; Robie et al., 2017). In automated video analysis of behavioral phenotypes of excitability mutants, however, identifying mutant-specific aberrant activities remains a challenge, as these behaviors are often complex motor sequences with substantially variable manifestations across individual flies.

In this report, we describe an approach to quantify aberrant movement events in flies carrying the *Shudderer* (*Shu*) allele based on automated video tracking locomotion data. *Shu* is a semi-dominant gain of function point mutation of the X-linked voltage-gated Na^+^ gene *paralytic* (i.e. *para*^*Shudderer*^, Kaas et al., 2016). Both female heterozygous *Shu/+* and male hemizygous *Shu/*Y flies display sporadic leg jerks and spontaneous “shuddering” convulsions, although phenotype is stronger in male flies (Williamson, 1982). These motor phenotypes are exacerbated at high temperatures (37 – 40 °C), with most *Shu/+* or *Shu/*Y flies eventually displaying convulsions and little other movement. Electrophysiological recordings from intact tethered flies reveal spontaneous spike bursts associated with seizures in *Shu/*Y flies, while *Shu/+* flies have a lower stimulation threshold for electroconvulsive seizure induction (Kaas et al., 2016; Kasuya et al., 2019).

We observed the behavior of *Shu* flies in an open-field arena placed on a temperature-controlled stage using a webcam (Landaverde et al., 2024). Initially, the stage temperature was set to a baseline temperature of ∼22 °C. After 180 s, the temperature gradually increased to ∼40 °C. Fly positions were tracked by IowaFLI Tracker, with each fly centroid represented as a [*x,y*] coordinate pair in each frame (Iyengar et al., 2012). Consistent with previous reports, *Shu* mutants at baseline temperatures displayed spontaneous shuddering events along with prolonged periods of ataxia consisting of immobilization with little movement (**Figure 1A**). Control flies (+/+ females and +/Y males) rarely showed these behaviors. Quantification of walking behavior indicated *Shu/+* mutants displayed reduced average velocities (*Shu/+*: 2.18 ± 0.09 vs. +/+: 5.91 ± 0.22 mm/s, mean ± SEM) and reduced path linearity (*Shu/+*: 0.30 ± 0.01 vs. +/+: 0.75 ± 0.02). *Shu/*Y mutants displayed similar differences in average velocity (*Shu/*Y: 2.14 ± 0.22 vs. +/Y: 6.55 ± 0.20 mm/s) and path linearity (*Shu/*Y: 0.30 ± 0.02 vs. +/Y: 0.80 ± 0.01). Also, in-line with previous water bath observations, at high temperatures (37 – 40 °C), we found *Shu/+* flies displayed more convulsions compared to the baseline temperature period but would continue to intermittently walk. *Shu/*Y flies appeared immobile at temperatures > 31 °C but showed occasional twitches. Control +/+ and +/Y flies continued to walk throughout the high-temperature period.

**Figure 1.**
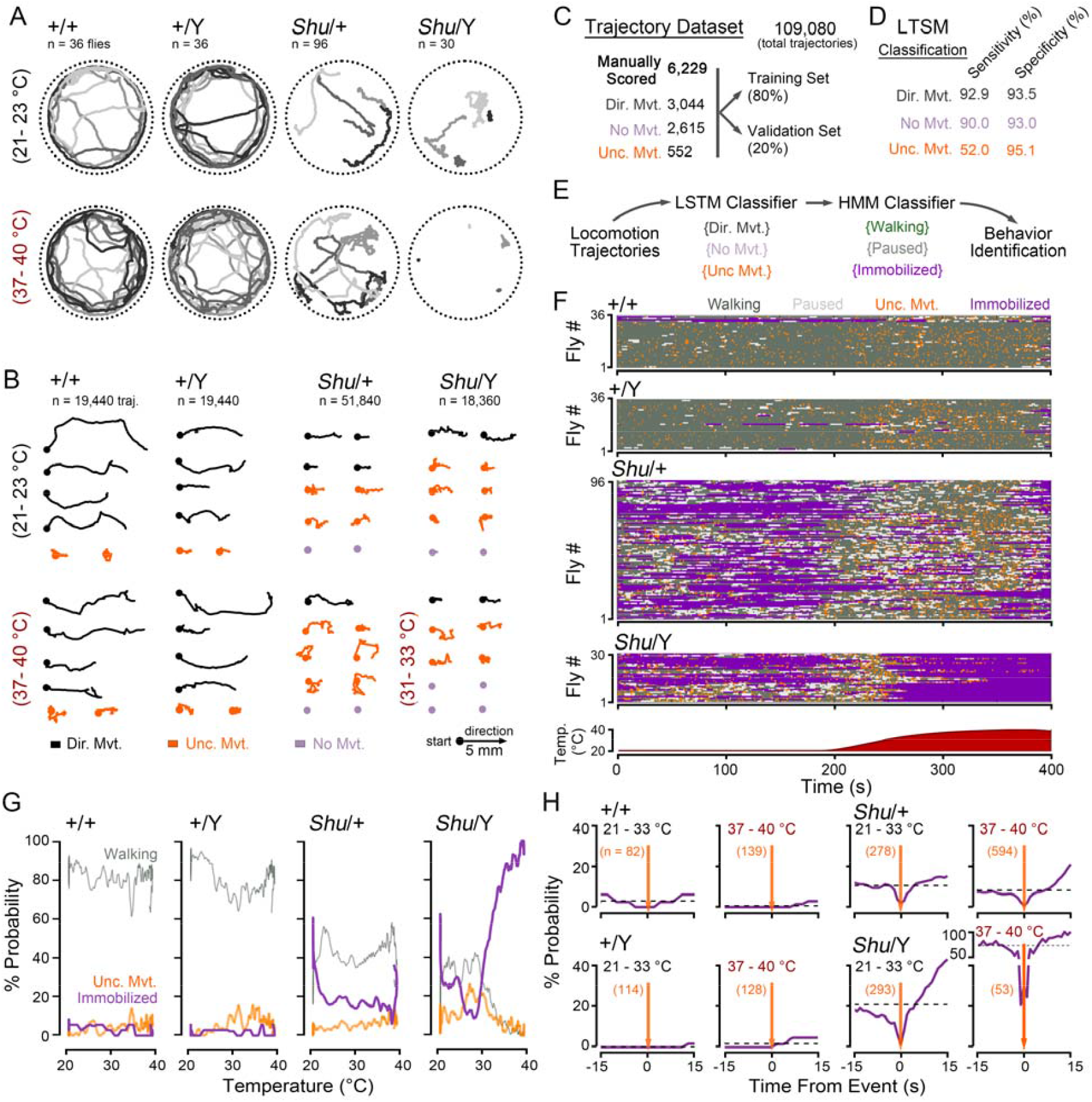
Automated classification of uncoordinated movements, walking, pauses, and immobilization in *Drosophila* wild-type flies and *Shu* mutants. (A) Representative fly movement tracks at baseline (21 – 23 °C) and high temperatures (37 – 40 °C) from the respective genotypes. Tracks from four flies are displayed; track duration: 20 s. (B) Example 2-s long movement trajectories. Trajectories are aligned to start at the dot and progress towards the right. Color indicates activity classification based on manual observer scoring (*directed movement, uncoordinated movement, no movement*). Total number of trajectories indicated. (C) Summary of the training data set. The number of trajectories manually classified in the respective events is indicated. Following manual classification, trajectories were randomly assigned to a “training” or “validation” dataset. (D) Performance of trained LSTM event classifier on the validation dataset. For each of the classified events, sensitivity is defined as the number of correctly classified events divided by the true total number of events; specificity is defined as the number of correctly classified non-events divided by the true total number of non-events. (E) Overall behavioral state classifier. Based on the time-series of LSTM events as an observed variable, a hidden Markov model trained to classify the fly as *walking, paused* or *immobilized*. (F) Raster plots of most likely activity state as classified by the hidden Markov model. Each row represents activity of a single fly over time (fly # indicated on y-axis). The corresponding stage temperature is indicated in the inset graph below. (G) Plots of the probability of observing activity states as a function of temperature. Sample size as in panel F. (H) Plots of the *immobilization* probability before and/or after an uncoordinated movement event (t = 0, orange line). Activity during the temperature ramp phase (21 – 33 °C) and high-temperature phase (37 – 40 °C) is plotted. Sample sizes indicate number of *uncoordinated movement* events. Dashed lines (black) indicate average expected probability of *immobilization*.

We sought to identify and quantify the spontaneous shuddering events and prolonged immobilization periods displayed by *Shu* mutants based solely on video tracking of fly centroids. Our overall strategy was to create a manually annotated dataset of normal and aberrant fly tracks amenable to train machine learning classifiers. Based on this training dataset, we could then create a system that classifies fly behavior at a particular moment based on the observed movement tracks. As shown in **Figure 1B**, the tracks from *Shu* mutants and control flies were segmented into 2-s long trajectories. A randomly selected subset of ∼ 6200 trajectories were each manually scored by an observer (blinded to the genotype) as either “*directed movement*”, “*no movement*” or “*uncoordinated movement*” (**Figure 1B-C**). Tracks categorized as *directed movement* corresponded with walking in both control flies and *Shu* mutants. In *Shu* mutants, *uncoordinated movements* events often indicated shuddering events, while in control flies these events corresponded with flies bumping into each other or falling from the lid. To enable robust development of classifiers based on the manually annotated dataset, we randomly split the dataset into a subset for training classifiers (∼80% tracks) and a subset intended for performance validation (∼20% tracks).

Using the manually annotated dataset, we trained an long short-term memory (LSTM) network (Hochreiter & Schmidhuber, 1997), a type of recurrent neural network suitable for analyzing time-series data, to classify unknown trajectories into one of the three movement categories.

Training of the LSTM was accomplished in ∼ 20 min using a desktop PC running MATLAB. Based on the validation dataset, the trained LSTM network was effective in classifying trajectories as *directed movement* or *no movement*, with both sensitivity (true positives rate) and specificity (true negative rate) greater than 90% (**Figure 1D**). For *uncoordinated movements* trajectories, we found the sensitivity to be lower (∼ 52%), but with a high specificity rate (95%). Thus, the *uncoordinated movements* identified by the LSTM reliably agreed with manual observation, but manually annotated events were sometimes missed by the LSTM.

In control flies, *no movement* trajectories appeared intermittently during brief (< 5 s) pauses in activity. In addition to these short duration pauses, *Shu* mutants also displayed prolonged immobilization events (> 10 s) consisting of nearly continuous sequences of *no movement* trajectories. To distinguish between the brief pauses and the immobilization events, we employed a hidden Markov model. In the model, the directly observed variable was the time series of LSTM classifications (i.e. *directed movements, no movements*, and *uncoordinated movements*) and we considered the fly to be in one of three states: *walking –* consisting of mostly *directed movements* trajectories; *paused –* consisting of mostly *no movement* trajectories with a high transition probability to the *walking* state; and *immobilized –* consisting of mostly *no movement* trajectories with a low transition probability to the *walking* state. Based on the training dataset, the HMM model generally classified sequences of *no movement* longer than ∼ 8 s as *immobilization*, while shorter sequences were classified as *pause* events. Thus, the overall classifier consisted of two stages: the LSTM that indicated *uncoordinated movements*, while the HMM classified *walking, pauses* and *immobilization* (**Figure 1E**).

The combined LSTM-HMM classifier was applied to the entire dataset of *Shu* and control fly data. To visualize classified fly activity patterns, we generated “actograms”— raster plots with each color signifying a particular activity, and each horizontal line depicting the time-series of a single fly’s activity during the temperature ramp protocol (**Figure 1F**). In the generated actograms, genotype-based differences are readily apparent. At the start of the ramp phase (21 – 23 °C), control flies displayed ongoing *walking* activity with occasional *pause* or *uncoordinated movement* events (**Figure 1G**). In these flies, *immobilization* was rarely observed (+/+: 5% of time, displayed by 2/36 flies, +/Y: 2% of time, displayed by 1/36 flies). In contrast, *Shu* mutants showed substantially more *immobilization*, with events observed in many *Shu/+* (34% of time, 65/100 flies) and *Shu/Y* flies (32% of time, 19/30 flies). In *Shu* flies, we found a high number of *uncoordinated movement* events despite the greatly reduced *walking* periods, suggesting these events are not related to general activity levels. Indeed, after removing *immobilization* events, in *Shu/+* flies *uncoordinated movement* events comprised 13% of all activity (compared to 5% in +/+ controls) flies. This fraction increased in *Shu/Y* flies, with *uncoordinated movement* accounting for 31% of all activity (vs. 4% in controls).

In the high-temperature period (37 – 40 °C), control flies continued to display normal walking activity with higher rates of *uncoordinated movement* events (+/+: 10%, +/Y: 9% of time) and an insignificant number of *immobilization* events (+/+: 1%, +/Y: 1% of time). *Shu* mutants, in contrast, displayed marked qualitative differences at high temperature. *Shu/+* mutants repeatedly transitioned between activity states, with most flies displaying *immobilization* at some point (82/100 flies). Excluding *immobilization* during this period, *uncoordinated movements* accounted for an increased proportion of activity (17%) in *Shu/+* flies. Male *Shu/*Y flies displayed a more extreme form of temperature sensitivity, with all flies displaying *immobilization* at the 37 – 40 °C temperature range (93% of time), occasionally punctuated by bouts of *uncoordinated movement* (5% of time). At high temperatures in these flies, no *walking* was detected. During the ramp protocol, the median temperature for *immobilization* event initiation was 31.7 °C (29.8 - 33.6 °C inter-quartile range).

Although the behavior classifier indicated *uncoordinated movement* events in both control and *Shu* mutants, we noted a key difference between events in the respective groups. In control flies, *walking* resumed immediately after an *uncoordinated movement* event, while *Shu* mutants in many cases did not recover *walking* activity (**Figure 1F**). These *uncoordinated movement – walking* events largely arose from flies falling from the arena ceiling or during social interaction with other flies. In contrast, *uncoordinated movement* events in *Shu* mutants often immediately preceded bouts of *immobilization* or occurred during periods of *immobilization* (**Figure 1H**).

Qualitative observation of *Shu* mutant videos indicated shuddering behaviors often corresponded with the *uncoordinated movement – immobilization* events. Indeed, the rate of these events was markedly higher in *Shu/+* flies (0.34 ± 0.07 min^-1^) compared to controls (0.06 ± 0.03 min^-1^). In *Shu/*Y males, the rate was even higher (1.29 ± 0.20 min^-1^) while the events were not observed in control +/Y flies (i.e. 0.00 ± 0.00 min^-1^).

This study aimed to develop a machine-learning system to quantify intermittent ataxic and convulsive movement events in a hyperexcitable *Drosophila* mutant *Shu* (Kaas et al., 2016; Williamson, 1982) based solely on video tracking data. Previous approaches to scoring aberrant hyperexcitable behaviors in *Drosophila* excitability mutants largely center on manual observation of induced behaviors, such as recovery from bang-sensitivity (Burg & Wu, 2012; Pavlidis et al., 1994) or sensitivity to high temperature (Palladino et al., 2003; Sun et al., 2012). Compared to manual human scoring, advantages of automated video tracking systems include a high experimental throughput, and consistent scoring performance. Our approach in developing such a system consisted of first creating a manually annotated dataset of movement trajectories from control flies and *Shu* mutants which was suitable for supervised training of machine-learning classifiers. Next, using the annotated dataset, an LSTM network was trained to classify movement trajectories (**Figure 1B**). We found a high level of LSTM performance in categorizing trajectories of walking-related *directed movement* and periods of *no movement* compared to manual scoring. For *uncoordinated movements*, the LSTM displayed moderate sensitivity but a high degree of specificity, with manually annotated events occasionally missed, but nearly all LSTM-indicated events in agreement with manual observation (**Figure 1C**). The mismatch between LSTM specificity and sensitivity may arise due to a relatively small training dataset or to constraints based on LSTM network design. Alternatively, intrinsic variability in manual classification could contribute to the phenomenon. Future studies utilizing larger dataset and efforts to systematically optimization classifier design will likely yield higher classification performance.

Next, using a previously developed strategy (Landaverde et al., 2024), we trained a HMM to determine one of three behavioral states: *walking, paused, immobilized*, based on the sequence of LSTM classifications of a fly’s movements. Results revealed clear distinctions in *Shu* mutant behavioral phenotypes, including bouts of *uncoordinated movements* followed by *immobilization* that were characteristic of qualitatively described shuddering behaviors (Kaas et al., 2016).

The expression of behavioral phenotypes in *Shu* mutants is remarkably variable. Previous reports indicate dietary factors including milk whey (Kasuya et al., 2019) and the ω-3 poly unsaturated fatty acid α-linolenic acid (Dinnel et al., 2025; Kasuya et al., 2023) as well as genetic background variation (Chen et al., 2020) can alter presentation seizure-related behavior. Our behavioral classifier enables a rapid, accessible and rigorous readout of genetic and environmental factors that modify *Shu* behavioral patterns. We also anticipate that with appropriate manually annotated training datasets, the classifier system may be extended to the large collection of fly mutants that display convulsions, ataxia or other episodic movement phenotypes. Many such mutants serve as models of neurological disease in humans (Bereshneh et al., 2025; Chi et al., 2022; Ehaideb et al., 2014; Hope et al., 2017; King et al., 2020; Marcogliese et al., 2022), and video tracking-based high-throughput behavioral characterization can facilitate neurogenetic analysis of underlying pathophysiology or screens for drug compounds that attenuate expression of behavioral phenotypes.

## Materials and Methods

### Fly stocks

The *paralytic*^*Shudderer*^ line (referred to as *Shu*) has previously been described in Kaas et al. (2016). The mutant strain is kept with an *FM7 w*^*-*^ balancer (i.e. *Shu/ FM7 w*^*-*^ *Bar*) and has previously been back-crossed to the *Canton-S* line for at least 10 generations. The *Canton-S* line is used as a control (+/+ or +/Y) line. Female heterozygous (*Shu/+*) and male hemizygous (*Shu/*Y) flies were generated by crossing *Shu/ FM7 w*^*-*^ *Bar* virgin females with male +/Y flies and collecting the resulting normal-eyed (non-*Bar*) progeny.

All flies were reared on a standardized fly media recipe used by the Bloomington *Drosophila* Stock Center. The recipe consisted of (per 1.0 l food): 918 ml H_2_O, 15.8 g yeast, 9.17 g soy flour, 67.1 g yellow cornmeal, 5.29 g agar, 71 ml corn syrup, 4.4 μl propionic acid. Flies were kept in a 25 °C incubator at 50% humidity and 12 h: 12 h light-dark cycle. For experiments, flies were collected within 24 h, and were aged 3 – 5 d.

### Video recording

The video recording procedure is based on Landaverde et al. (Landaverde et al., 2024), using IowaFLI Tracker for fly tracking (Iyengar et al., 2012). Four flies were loaded into each of four circular behavioral arenas (dia. 28 mm) in the video tracking system (16 flies total per video). The arenas were seated on Whatman #1 filter paper and placed on a Peltier temperature-controlled stage (AHP 1200 CPV, TECA Corp). A T-type thermocouple connected to a data acquisition card (TC-01, National Instruments) monitored arena temperature, and a custom-written LabView program controlled stage temperature.

Fly behavior was monitored by a webcam (c920 Logitech) recording at 30 frames per second controlled by the LabView program. The fly positions were tracked by IowaFLI tracker, yielding a time-series of positional *x*_*n*_ and *y*_*n*_ coordinates in each frame (*n*) for each fly’s path. Fly velocity, % time active and distance traveled were calculated as described in (Landaverde et al., 2024).

For each second of video, a fly trajectory was generated. Each trajectory consisted of 60 frames (2 s) of positional data 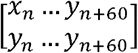. The fly trajectory corresponding with frame *n* of video was computed as follows:

1. Center the fly position 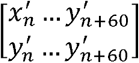 by subtracting the initial position coordinates:

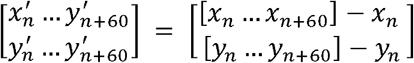
2. Rotate the trajectory 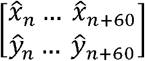 such that the last coordinates (*x*_*n+60*_, *y*_*n+60*_) are aligned on the x-axis, (i.e. faces rightwards). The rotation angle *θ* is: *θ* = atan2(*y*′_*n*+60_, *x*′_*n*+60_).

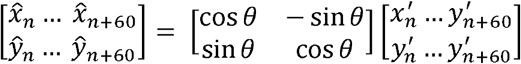

### Manual observer classification

A random selection of fly trajectories (n = 6,229) was classified manually into one of three categories: “directed movement”, “no movement” or “uncoordinated movement”. A MATLAB script generated a plot of randomly selected fly trajectories and a human observer scored the trajectory as one of the three categories. Observers were blinded to the genotype, temperature and elapsed time corresponding with the trajectory.

### LSTM classifier training

Training of the long short-term memory (LSTM) network on the manually observed fly trajectory data set was done in MATLAB (Mathworks r2024b) using the Deep Learning Toolbox. The training dataset consisted of ∼80% manual classified trajectories. The LSTM consisted of 180 bidirectional hidden layers implemented by the ‘*bilstmLayer’* function. Cross-entropy was used as the loss function, with class weights normalized to the relative frequency of “directed movement”, “no movement” and “uncoordinated movement” classifications in the training dataset. LSTM training was done using the ‘*trainnet’* function in MATLAB, using the ‘*adam’* solver, ‘*max epoch*’ = 200, ‘*initial learn rate*’ of 0.002 and ‘*gradient threshold’* of 1. All other Deep Learning Toolbox parameters were set to default values.

Classification of trajectories using the LSTM was done by the ‘*minibatchpredict*’ function. For each input trajectory, the LSTM returned a set of three scores corresponding to the likelihood of a “directed movement”, “no movement” or “uncoordinated movement” label.

### Hidden Markov Model

To classify immobilization, pausing and walking activity states, a hidden Markov model (HMM) was trained using the LSTM classification of “directed movement” vs. not “directed movement” time-series as an observed variable. The HMM had three hidden states, “walking”, “paused” (short non-walking events), and “immobilized” (long non-walking events). The HMM was trained in MATLAB using the ‘*hmmtrain’* function. The trained estimated transition matrix (**TRANS**) was:

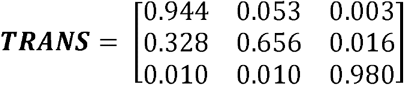

The trained emission matrix (**EMIS**) was:

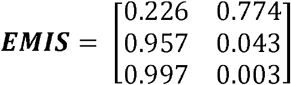

For a given time-series of “directed movement” versus not “directed movement”, the ‘*hmmdecode*’ function generated a likelihood of “walking”, “paused”, or “immobilized” activity states.

### Machine learning implementation and statistical analysis

All machine learning and statistical analysis was done in MATLAB (r2024b, MathWorks). A Dell XPS PC running an Intel i9-12900K processor with 64 GB RAM and an AMD 6700 XT graphics card was used for all analysis. Sample sizes are indicated in figure legend.

## Author Contribution

AI and VH designed behavioral experiments; VH produced the behavioral analysis dataset; AI & VH developed new software and analysis methodology; AI wrote MATLAB code; AI drafted the initial manuscript; AI and VH edited and approved final manuscript draft.

## Acknowledgments

We thank Reid Schuback, Aubrey Gray, Bailey O’Neal and other Iyengar Lab members for their technical assistance. We appreciate Toshi Kitamoto’s and Lawrence Reiter’s critical comments and advice during this project. This work was supported by a UA FUSE Award and NIH Grant (NS 134960) to AI, and UA ASSURE Award to VH.

